# Construction and optimization of a heterologous pathway for protocatechuate catabolism in *Escherichia coli* enables rapid bioconversion of model lignin monomers

**DOI:** 10.1101/094805

**Authors:** Sonya M. Clarkson, Donna M. Kridelbaugh, James G. Elkins, Adam M. Guss, Joshua K. Michener

**Author notes:** Correspondence should be sent to JKM or AMG. This manuscript has been authored by UT-Battelle, LLC under Contract No. DE-AC05-00OR22725 with the U.S. Department of Energy. The United States Government retains and the publisher, by accepting the article for publication, acknowledges that the United States Government retains a non-exclusive, paid-up, irrevocable, world-wide license to publish or reproduce the published form of this manuscript, or allow others to do so, for United States Government purposes. The Department of Energy will provide public access to these results of federally sponsored research in accordance with the DOE Public Access Plan (http://energy.gov/downloads/doe-public-access-plan).

## Abstract

Cellulosic biofuel production yields a substantial lignin byproduct stream that currently has few applications. Biological conversion of lignin compounds into chemicals and fuels has the potential to improve the economics of cellulosic biofuels, but few microbes are able both to catabolize lignin and generate valuable products. While *Escherichia coli* has been engineered to produce a variety of fuels and chemicals, it is incapable of catabolizing most aromatic compounds. Therefore, we have engineered *E. coli* to catabolize a model lignin monomer, protocatechuate, as the sole source of carbon and energy, via heterologous expression of a nine-gene pathway from *Pseudomonas putida* KT2440. We next used experimental evolution to select for mutations that increased growth with PCA more than two-fold. Increasing the strength of a single ribosome binding site in the heterologous pathway was sufficient to recapitulate the increased growth. After optimization of the core pathway, we extended the pathway to enable catabolism of a second model compound, 4-hydroxybenzoate. These engineered strains will be useful platforms to discover, characterize, and optimize pathways for lignin bioconversions.

**Highlights:**

- A heterologous pathway for PCA catabolism was transferred to Escherichia coli.
- Evolution identified a mutation that increased growth with PCA by 2.5-fold.
- Optimization plus further engineering allowed efficient catabolism of 4-HB

## 1. Introduction

Cellulosic biofuels will likely play an important role in the transition to a sustainable and carbon-neutral economy (Department of Energy, 2016). In a typical biotransformation, the carbohydrate-rich cellulose and hemicellulose are extracted and fermented to yield the desired biofuel. The lignin, comprising more than 25% of the dry biomass, is generally then burned or diverted to other low-value uses (Ragauskas et al., 2014). Lignin is a challenging feedstock for chemical or biochemical transformations due to its inherent heterogeneity, which varies widely depending on the biomass source and pretreatment strategy. Developing new high-value uses for lignin will improve the economics of cellulosic biofuels.

Despite the challenges involved in degrading lignin, its ubiquity in nature means that microbes have evolved to catabolize it. White-rot fungi were the first lignin-degrading microbes to be extensively described, and numerous lignin-degrading bacteria have also been isolated and characterized (Bugg et al., 2011). Using promiscuous oxidative enzymes to depolymerize lignin and specific pathways to catabolize the resulting products, microbes are capable of converting complex lignin-rich feedstocks into valuable bioproducts (Linger et al., 2014; Salvachúa et al., 2015). However, no microbe has yet been identified that is sufficiently effective at both converting lignin into common metabolic intermediates and converting those intermediates into useful bioproducts at high titer, rate, and yield. Engineering and characterizing lignin degradation pathways in model organisms would provide an opportunity to combine these two characteristics.

*Escherichia coli* is a major platform organism for metabolic engineering and synthetic biology due to the availability of extensive genetic, biochemical, and physiological information and the relative ease of genetic modification. *E. coli* is routinely used to prototype metabolic pathways for the production of fuels and chemicals [add miscellaneous references]. However, *E. coli* is incapable of using lignin-derived aromatic compounds as a carbon and energy source, limiting its utility for the bioconversion of the lignin fraction of lignocellulose.

For many organisms that are capable of catabolizing aromatic compounds, protocatechuic acid (PCA) is a key metabolic intermediate in this pathway. More complex lignin compounds are metabolized first into PCA before being assimilated into central metabolism (Jimenez et al., 2002). Three pathways for degrading PCA have been characterized, differing in the location of the initial ring-opening oxidation. Representatives from each class of pathway have been expressed from plasmids in *E. coli*, with varying levels of success. The meta-cleavage pathway has been best characterized in *Sphingobium* sp. SYK-6 (Masai et al., 2007), and heterologous expression in *E. coli* of a pathway variant from *Pseudomonas ochraceae* NGJ1 allowed colony formation after two days (Maruyama et al., 2004). The para-cleavage pathway has been studied in *Paenibacillus* sp. JJ-1b, and heterologous expression of this pathway in *E. coli* enabled colony formation after a ten-day incubation (Kasai et al., 2009). Finally, the most widely distributed pathway begins through ortho-cleavage, as typified by *Pseudomonas putida* (Harwood and Parales, 1996; Ornston and Stanier, 1964) (Figure 1A). Expression of the ortho-cleavage pathway from *Acinetobacter calcoaceticus* in *E. coli* quantitatively produced an intermediate, (β-ketoadipate, but did not allow growth (Doten et al., 1987).

**Figure 1:**
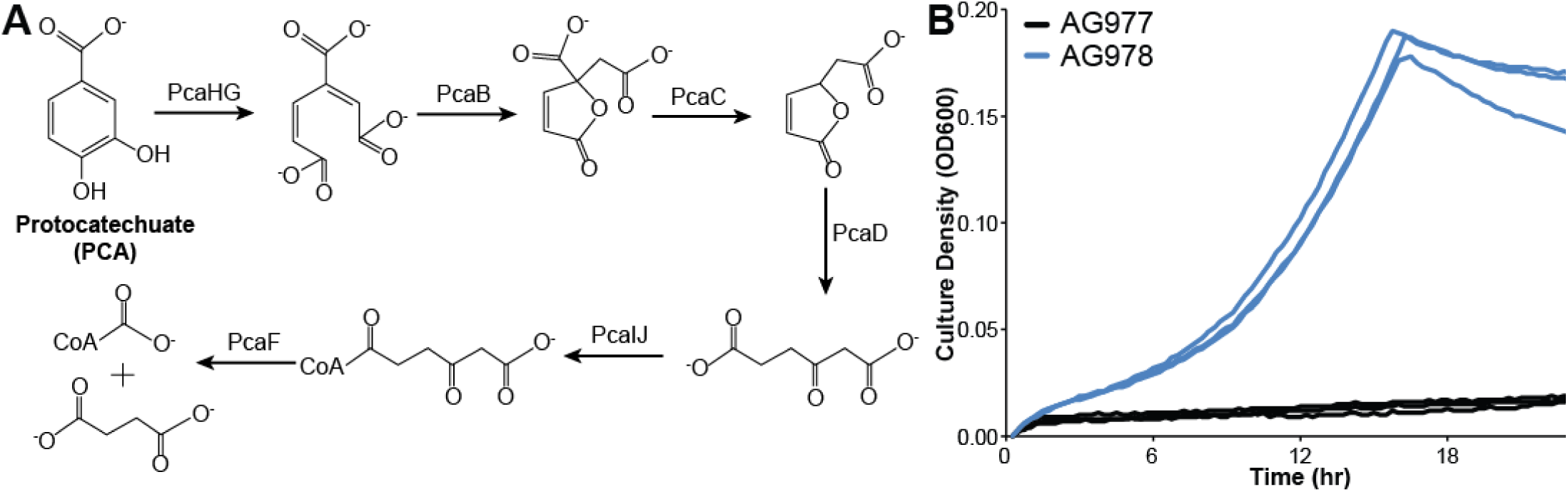
(A) The pathway for ortho-cleavage of PCA was transferred from *P. putida* KT2440 to *E. coli*, allowing growth with PCA as the sole source of carbon and energy. (B) Strain AG978 (*∆ompT::pcaHGBDC ∆pflB::pcaIJFK)* was grown in minimal medium containing PCA as the sole source of carbon and energy. The control strain, AG977 (*∆ompT ∆pflB*) shows a slow increase in absorbance due to oxidation of the substrate.

When microbes grow poorly on a particular substrate, experimental evolution can be used to optimize inefficient metabolic pathways, whether native (Herring et al., 2006; Hong et al., 2011) or engineered (Chou et al., 2011; Clark et al., 2015; Michener et al., 2014). Serial propagation under conditions where the relevant pathway is necessary for growth selects for mutants with improved pathway function. Genome resequencing can then be used to identify putative causal mutations. Reconstructing these mutations in the parental strain verifies their effects. By studying the mutations that improve pathway function, we can deduce the factors that were initially limiting pathway effectiveness and the biochemical solutions that overcame these limitations.

In this work, we have combined rational and evolutionary approaches to construct and optimize a heterologous pathway for PCA catabolism in *E. coli.* We first transferred the ortho-cleavage pathway from *P. putida* KT2440 into E. *coli* and demonstrated that the engineered strain can grow with PCA as the sole source of carbon and energy. Next, we used experimental evolution to identify a single mutation that increased the growth rate with PCA while maintaining basal growth rates with glucose. Finally, we showed that the optimized PCA degradation pathway enables further extension of the catabolic network to the related compound 4-hydrobenzoate (4-HB). This optimized strain will serve as a platform for further reconstruction of lignin catabolic pathways.

## 2. Materials and Methods

### 2.1 Media and chemicals

All chemicals were purchased from Sigma-Aldrich (St. Louis, MO) or Fisher Scientific (Fairlawn, NJ) and were molecular grade. All oligonucleotides were ordered from IDT (Coralville, IA). *E. coli* strains were routinely cultivated at 37 °C in LB broth containing the necessary antibiotics (50 mg/L kanamycin, 50 mg/L carbenicillin, 15 mg/L chloramphenicol, or 50 mg/L spectinomycin). Growth assays with PCA and 4-HB were performed in M9 salts medium containing 300 mg/L thiamine and 1 mM isopropyl β-D-1-thiogalactopyranoside (IPTG). PCA and 4-HB were dissolved in water at 5 g/L, filter sterilized, and added at a final concentration of 1 g/L. The pH of the substrates was not controlled, as PCA oxidation occurred more rapidly at neutral pH.

### 2.2 Plasmid construction

The eight genes of the PCA ortho-degradation pathway from *Pseudomonas putida* KT2440, *pcaHGBCDIJF,* and the PCA transporter, *pcaK*, were designed as four constructs using GeneDesigner software and ordered through DNA2.0 (Menlo Park, CA). The genes were codon optimized for *E. coli*, placed behind an inducible T5-*lac* promoter, and flanked by BioBrick restriction enzyme sites for cloning. Constructs included ribosome binding sites and re-initiation spacer sequences between genes.

Plasmids for chromosomal integration of the PCA ortho-degradation pathway were constructed in pET30a (EMD Millipore, Billerica, MA). The plasmid pET30a-dest was created by blunt-end cloning the PCR product created with primers dest-FWD / dest-REV into pET30a. Ligation products were transformed into CopyCutter EPI400 *E. coli* (Epicentre, Madison, WI) and plasmids were verified by PCR with primers T7 promoter / T7 terminator. The final plasmids used for integration were constructed in pET30a-dest using standard methods to insert PCR products generated using I-SceI-LP-T5-FWD and LP-IJFK-REV, LP-HGBDC-REV, or LP-control REV, resulting in pET30a-LP-control, pET30a-LP-pcaIJFK, and pET30a-LP-pcaHGBDC.

A plasmid containing both the *λ* Red recombinase system and the homing endonuclease I-*sce*I was used for chromosomal insertion of the PCA ortho-degradation pathway. The I-*sce*I gene was codon-optimized for *E. coli*, placed behind a strong ribosome binding sequence (Life Technologies, Grand Island, NY), and inserted upstream of the *λ* Red recombinase genes on plasmid pKD46 using standard methods (Datsenko and Wanner, 2000), resulting in the plasmid pKD46-I-SceI. I-SceI enzyme functionality was tested in an overnight culture grown with 1 mM arabinose to induce I-SceI expression. Cells were harvested, washed, and frozen at −20°C before lysis in I-SceI buffer (10 mM TrisHCl, 10 mM MgCl2, 1 mM DTT, 100 mg/mL bovine serum albumin, pH 8.8). Cell extract was incubated with the plasmid pGPS2 (New England Biolabs, Waltham, MA) at 37°C for 1 hour. Samples were taken every 15 minutes and analyzed by gel electrophoresis to confirm I-SceI digestion activity (Figure S1B).

Plasmids pJM180 and pJM182, expressing tracrRNAs targeting the *pcaH* RBS were generated through inverse PCR of pTarget using primers pTarget FWD and pTarget REV (Jiang et al., 2015). The appropriate oligos were then assembled into the pTarget backbone using the HiFi master mix and transformed into 10-β *E. coli* (NEB, Waltham, MA). Plasmid pJM179, targeting *flu*, and plasmids pJM160, targeting *elfC* for integration of *praI*, were constructed in a similar fashion.

### 2.3 Strain construction

The PCA ortho-degradation pathway was inserted into the chromosome of *E. coli* BW25113 using a “Landing Pad” method modified from Kuhlman and Cox (2010). A chloramphenicol resistance gene cassette flanked by I-SceI restriction sites and “landing pad” regions of homology for *λ* Red recombination (1: 5’-TACGGCCCCAAGGTCCAAACGGTGA-3’ and 2: 5’-GATGGCGCCTCATCCCTGAAG CCAA-3’) was PCR amplified from pUC57-LP with primers ompT-LP-up / ompT-LP-down or pflB-LP-up / pflB-LP-down to add flanking 50 bp homology regions up- and down-stream of either *ompT*or *pflB*. The ompT PCR product was transformed into *E. coli* BW25113 containing pKD46 and expressing the λ Red recombinase system. Chloramphenicol-resistant colonies were screened for gene replacement by PCR at *ompT* (ompT-FWD / ompT-REV). Correct strains were co-transformed with pKD46-I-SceI and either the pET30a-LP-control or pET30a-LP-pcaHGBDC plasmid and selected using kanamycin and carbenicillin. Plasmid transformation was confirmed by restriction digest and strains containing both plasmids were grown in LB + Kan + Carb at 30°C for 3 h, harvested, re-suspended in LB + Carb with 1 mM arabinose to induce the λ Red recombinase system and I-SceI expression, and grown at 30°C for 1 h. Culture (10 μL) was streaked on LB plates and incubated at 37°C overnight to cure the pKD46-I-SceI plasmid. Colonies were patched to LB and LB + Cm at 37°C and all Cm-sensitive colonies were PCR screened at *ompT* Correct strains for each integration (∆*ompT* and *ompT::pcaHGBDC*) were streaked on LB, patched on LB, LB+Kan, and LB+Carb at 30°C to verify expected phenotypes, and all Kan- and Carb-sensitive colonies were again PCR screened at *ompT* to confirm integration. The process was repeated in the *∆ompT* and *∆ompT::pcaHGBDC* strains for *∆pflB* or replacement with *pcaIJFK,* resulting in *∆ompTΔpflB* (AG977) and *∆ompT::pcaHGBDC ∆pflB::pcaIJFK*(AG978). Strains correct by PCR were sequence verified at both the *ompT* and *pflB* loci.

### 2.4 Growth measurements

Cultures were grown to saturation in LB + 1 mM IPTG, then diluted 100-fold into fresh M9 + 1 mM IPTG + 0.2% glucose and regrown to saturation. Cultures were then diluted 100-fold into fresh M9 + 1 mM IPTG + substrate and grown as triplicate 100 μL cultures in a Bioscreen C plate reader (Oy Growth Curves Ab Ltd, Helsinki, Finland). Growth was monitored using optical density at 600 nm (OD_600_). Growth rates were calculated using CurveFitter software (Delaney et al., 2013).

### 2.5 Experimental evolution

The parental strain, AG978, was streaked to single colonies. Three separate colonies were grown to saturation in M9 + 1 mM IPTG + 0.2% glucose, and then diluted 128-fold into M9 + 1 mM IPTG + 1 g/L PCA. When the cultures reached saturation, initially after 48 hours but later shortened to 24 hours, they were diluted a further 128-fold into fresh M9 + 1 mM IPTG + 1 g/L PCA. Culture aliquots were periodically frozen at −80°C for later analysis. After 500 generations, each culture was streaked to single colonies. Eight colonies from each plate were picked and analyzed for growth in M9 + 1 mM IPTG + 1 g/L PCA.

### 2.6 Genome resequencing and reconstruction

Genomic DNA was prepared using a DNeasy Blood and Tissue kit (Qiagen, Valencia, CA) according to the manufacturer’s directions. Purified DNA was quantified using a Qubit fluorimeter (Thermo Fisher, Waltham, MA), resequenced by the Joint Genome Institute using a MiSeq (Illumina, San Diego, CA) to approximately 150x coverage, and analyzed using Geneious (Biomatters, Auckland, NZ). Targeted mutations were introduced into the parental strain on ssDNA oligonucleotides or dsDNA gBlocks using *λ*-RED in combination with CRISPR/Cas (Jiang et al., 2015). Mutations were verified by Sanger sequencing of PCR amplicons. The gene cassette for *elfC::praI* was synthesized by Gen9 (Cambridge, MA) and integrated into the appropriate chromosomal locus following the same process as for the targeted mutations. The promoter and terminator for *praI* expression were chosen from previously characterized genetic parts (Chen et al., 2013; Kosuri et al., 2013).

### 2.7 RBS design and analysis

The parental and mutant ribosome binding sites were analyzed using the RBS Calculator with Free Energy Model v2.0 (Espah Borujeni et al., 2014). Mutant ribosome binding sites were designed using the RBS Calculator (Salis et al., 2009).

### 2.8 qPCR

Gene copy numbers were determined by quantitative PCR using Phusion DNA Polymerase (NEB) and EvaGreen (Biotium, Fremont, CA) on a CFX96 Touch thermocycler (Bio-Rad, Hercules, CA). Whole cells from triplicate cultures were used as the DNA templates. *pcaH* was measured using primers pcaH_SeqF (5’-CGCTCACAATTCCACAACG-3’) and pcaH_SeqR (5’-CTTTTGGGTGCCAATTTCTATC-3’). Copy number was normalized to 16S rRNA gene copies determined using primers 515F (5’GTGCCAGCMGCCGCGGTAA-3’) and 1492R (5’-GGTTACCTTGTTACGACTT-3’).

## 3. Results and Discussion

### 3.1 PCA ortho-degradation pathway design and construction

The eight genes of the PCA ortho-degradation pathway from *Pseudomonas putida* KT2440, *pcaHGBCDIJF*, and the PCA transporter, *pcaK*, were codon optimized for *E. coli* and combined into two operons, each driven by an inducible T5-*lac* promoter. These operons were synthesized *de novo* and inserted into the chromosome of *E. coli* BW25113 at the *ompT* and *pflB* loci, yielding strain AG978 (*∆ompT::pcaHGBDC ∆pflB::pcaIJFK*).

Expression of the PCA pathway allowed growth with PCA as the sole source of carbon and energy, with a growth rate of 0.16 hr^-1^ (Figure 1B). Control strains lacking the pathway genes showed a slow increase in optical density due to the formation of a red color, likely caused by oxidation of PCA, but no increase in turbidity or cell count was observed.

### 3.2 Evolution improved function of the heterologous PCA pathway

While strain AG978 grows with PCA, its growth rate is roughly one quarter that of *P. putida* grown under similar conditions (Nichols and Harwood, 1997). To optimize this pathway, we used experimental evolution to select for mutants with improved growth on PCA. Three replicate cultures of AG978 were grown in minimal medium with 1 g/L PCA as the sole source of carbon and energy. When each culture reached saturation, it was diluted 128-fold into fresh medium and allowed to regrow. After 500 generations, individual mutants were isolated and characterized. Each of the evolved isolates grew significantly faster than the parent, with improvements between 2.2-fold and 2.4-fold (Figure 2). We identified mixed phenotypes in two of the evolved populations. Replicate population A yielded strains JME1 and JME2, replicate population B yielded strains JME3 and JME4, but all of the isolates from replicate population C showed the same phenotype as JME6. While all of the isolates had similarly improved growth rates, JME1, JME4, and JME6 reached a substantially lower final OD, even when compared to the unevolved strain.

**Figure 2:**
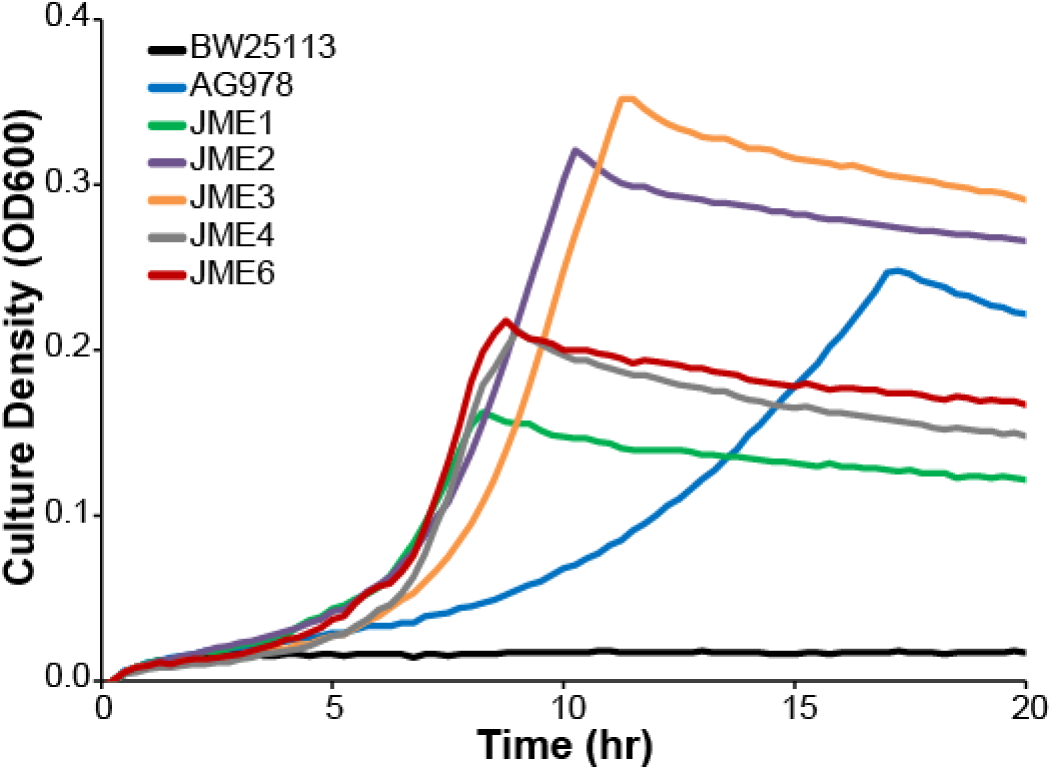
Experimental evolution selects for improved growth with PCA. Five strains, isolated from three replicate evolution experiments, were compared to the wildtype BW25113 and the engineered parent AG978 during growth in minimal medium containing 1 g/L PCA. Each of the evolved isolates grows significantly faster than the parent. Three of the isolates, JME1, JME4, and JME6 show a reduced maximum optical density.

### 3.3 Genome resequencing and reconstruction identify causal mutations

We resequenced the genomes of all five isolates to identify mutations relative to the parental strain. The isolates had between five and eight mutations, including several large IS-mediated deletions and rearrangements (Supplementary Table 4). Many strains shared independent mutations in the same genes or genomic regions, even across replicate cultures.

All five strains had mutations adjacent to the *pcaH* gene, which catalyzes the first step in PCA oxidation (Figure 1A and Supplementary Figure 1). Strains JME1, JME2, JME3 had IS-mediated duplications of a 118-kb region containing the introduced *pcaHGBDC* operon. Strain JME4 had a single nucleotide mutation in the predicted ribosomal binding site (RBS) for *pcaH* (Supplementary Figure 1). Using the RBS Calculator, the RBS mutation in JME4 is predicted to increase the strength of the *pcaH* RBS by 2.2-fold (Salis et al., 2009). Strain JME6 contains an IS element inserted between the promoter and RBS of *pcaH* that is likely to increase expression of the PcaH operon (Schnetz and Rak, 1992). In combination, these results suggest that increasing PcaH expression, either through changes in copy number, transcription rate, or translation rate, is highly beneficial during growth with PCA.

Reconstructing IS-mediated rearrangements is challenging, so to test this hypothesis we introduced the single nucleotide mutation from JME4 into the parental AG978 strain, yielding strain JME17. This single mutation recapitulated the majority of the evolutionary improvement in growth with PCA, raising the growth rate from 0.16 hr^-1^ to 0.41 hr^-1^ (Figure 3A). Comparing JME17 to the evolved isolate JME3, JME3 grows 24% faster with PCA and 20% faster with glucose (Figure 3B). Since JME3 displays a similar improvement in growth relative to JME17 when grown with either glucose or PCA, the unexamined mutations in JME3 likely reflect general adaptations to growth in minimal medium under these experimental conditions rather than any PCA-specific adaptation. Our results with PcaH are consistent with previous studies that have shown large changes in growth rate resulting from small changes in gene expression of metabolic enzymes (Kershner et al., 2016; Michener et al., 2014).

**Figure 3:**
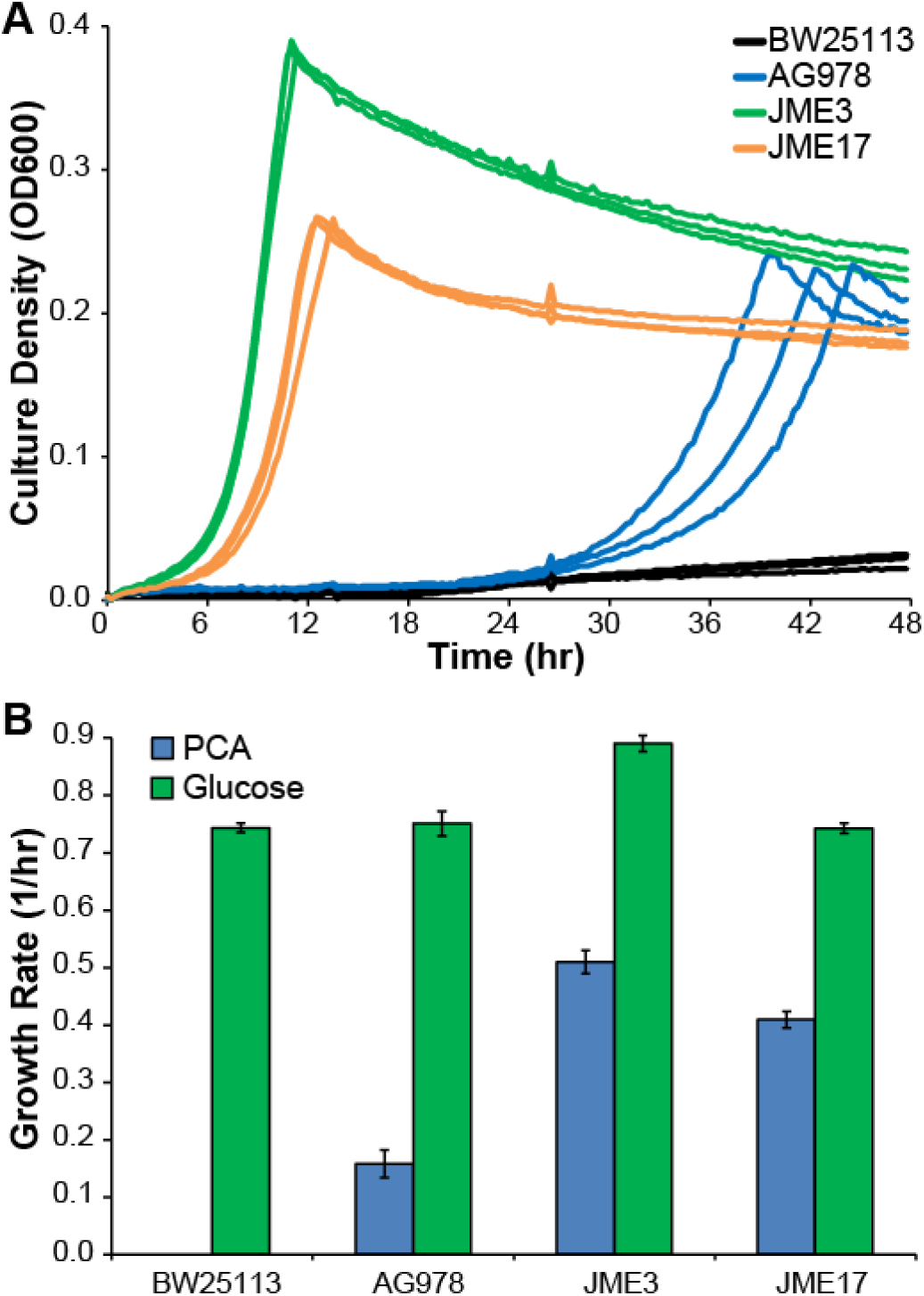
A mutation in the RBS of *pcaH* is sufficient to reproduce the increase in PCA-specific growth rate. (A) Comparing the growth of the evolved isolate JME3 with the reconstructed strain JME17 demonstrates that both strains grow more rapidly with PCA than the parent AG978. JME17 differs from AG978 by a single point mutation in the RBS of *pcaH*. (B) Comparing growth rates in minimal medium containing 1 g/L PCA or 2 g/L glucose, JME3 has a moderate growth advantage relative to JME17 in both conditions. Error bars show one standard deviation, calculated from three biological replicates.

Three of the strains, JME1, JME4, and JME6 reached saturation at relatively low optical densities (Figure 2). The only mutational target common to these three strains, but not to JME2 or JME3, is the surface antigen Ag43 encoded by *flu*. JME1 contains a nonsynonymous mutation in *flu*, while JME4 and JME5 have non-overlapping 156-nt and 261-nt deletions, respectively. We introduced the 261-nt deletion from JME6 into the parental strain and into the strain carrying the *pcaH* RBS mutation, yielding strains JME15 and JME16. In both cases, the *flu* deletion reduced the maximum optical density without significantly affecting growth rate (Supplementary Figure 2). The replicate evolution of *flu* mutations across all three populations strongly suggests that the mutation was beneficial, and a similar mutation has been seen in a previous evolution experiment looking at adaptation to acid stress (Harden et al., 2015). All of the mutations observed in this experiment produce modified forms of Ag43, rather than deletions or frameshift mutations. Ag43 mediates cell aggregation (Diderichsen, 1980), and modulation of aggregation by these mutant forms of Ag43 may increase cell survival under acid-stressed conditions such as minimal medium containing PCA.

### 3.4 Targeted RBS mutation replaced large-scale gene duplication

In replicate population B, isolates JME3 and JME4 demonstrate two different mechanisms to increase PcaH expression. JME3 contains a 118-kb duplication that spans the entire heterologous gene cluster containing *pcaH*. In contrast, JME4 has only a single copy of that region and, instead, has a single nucleotide mutation in the RBS of *pcaH*. Both the RBS mutation and tandem duplication would be predicted to increase PcaH expression by roughly two-fold. Targeted Sanger sequencing shows that the RBS mutation seen in JME4 rises to an observable frequency between generations 400 and 500 (data not shown). Conversely, qPCR results show that the population-averaged copy number of *pcaH* increases to more than three copies per cell before declining between generations 400 and 500 (Supplementary Figure 3A). In combination, these results demonstrate that the tandem duplication evolved first and was then in the process of being replaced by the RBS mutation between generations 400 and 500. Due to possible clonal interference, the replacement of the JME3 lineage with the JME4 lineage cannot be attributed solely to the different mechanisms for overexpressing PcaH since the JME4 lineage may have additional beneficial mutations. However, similar evolutionary trajectories of transient duplication and refinement have been observed in evolving populations of *E. coli* and *Saccharomyces cerevisiae* (Blount et al., 2012; Yona et al., 2012).

### 3.5 Additional increases in *pcaH* RBS strength do not improve growth

To further explore the importance of the *pcaH* RBS for rapid growth, we designed three additional ribosome binding sites with predicted 4-, 8-, and 16-fold increases in RBS strength relative to the parental strain (Salis et al., 2009). These strains showed similar growth rates to the engineered mutant JME17 (Supplementary Figure 3B). The two-fold difference between the predicted *pcaH* RBS strengths in AG978 and JME17 is similar to the potential uncertainty in the RBS Calculator prediction. However, the fold changes for the engineered RBSs are larger than the prediction uncertainty. Since all of the modified RBSs increase the growth rate relative to AG978, it is likely that all of the modified RBSs increase the RBS strength. Additionally, since the engineered RBSs do not increase the growth rate relative to JME17, the increase due to the single nucleotide mutation in JME17 is sufficient to relieve any growth restriction due to limited PcaH expression.

### 3.6 Improved PCA catabolism supports further pathway elaboration

A single enzyme, 4-hydroxybenzoate 3-monooxygenase, can convert 4-HB into PCA. This enzymatic activity can be provided by PraI from *Paenibacillus* sp. JJ-1B (Kasai et al., 2009). We introduced the corresponding gene into the parental strain AG978 and the engineered strain JME17, yielding strains JME7 and JME50, respectively (Figure 4A). When cultured with 4-HB as the sole source of carbon and energy, JME7 grew at a rate of 0.027 ± 0.005 hr^-1^, while JME50 grew at 0.259 ± 0.002 hr^-1^ (Figure 4B). The RBS mutation in *pcaH* is the only difference between JME7 and JME50; while this mutation increased growth with PCA by roughly 2.5-fold, it increased growth with 4-HB by nearly 10-fold. Further extension of the catabolic network to include new growth substrates will require careful optimization of each new pathway.

**Figure 4:**
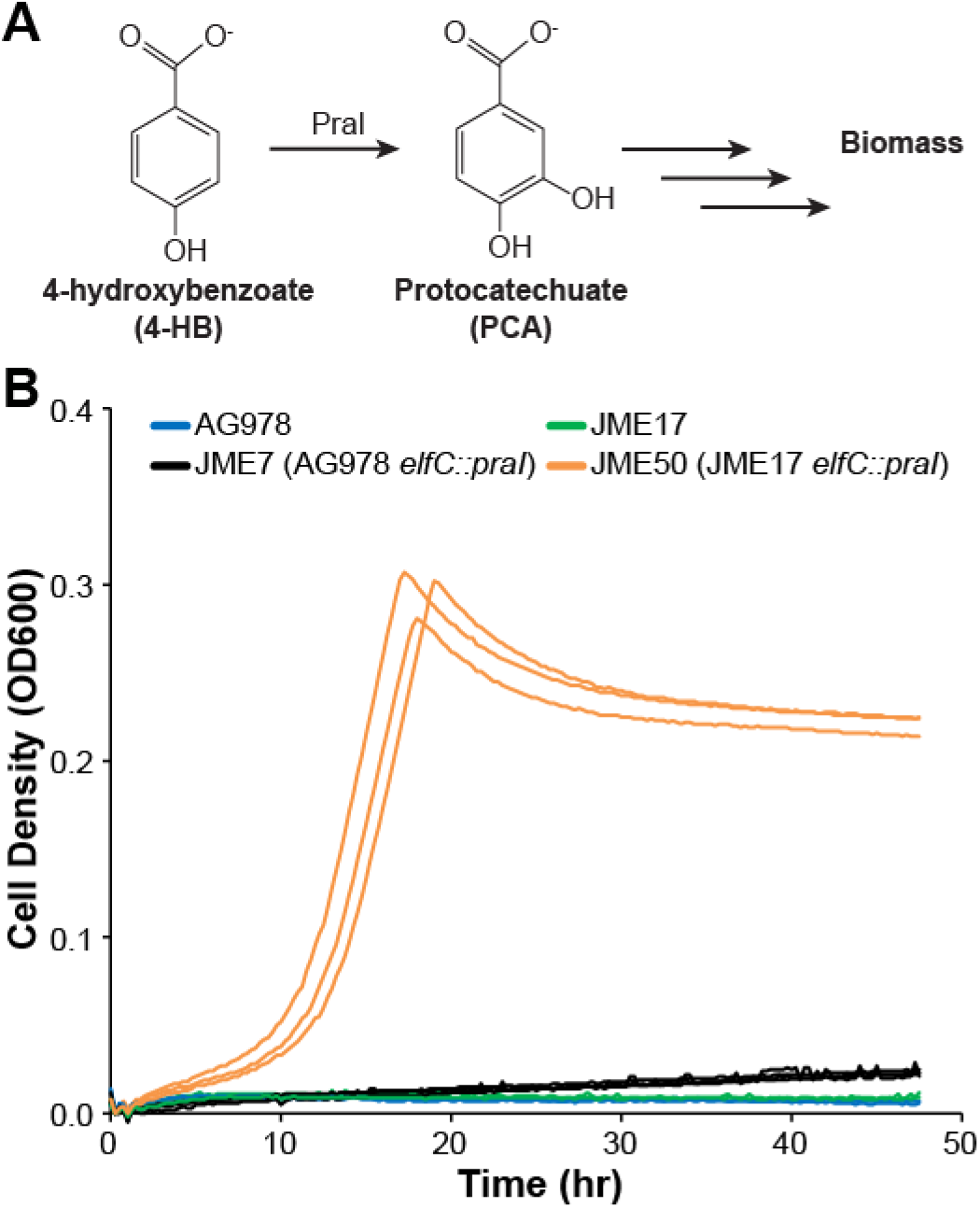
Evolutionary optimization of the core PCA catabolism pathway enables elaboration of the catabolic network. (A) The 4-hydroxybenzoate 3-monooxygenase Pral converts 4-HB into PCA. (B) The monooxygenase gene, *praI*, is integrated in both JME7 (black) and JME50 (orange). The difference between JME7 and JME50 is a single point mutation in the RBS of *pcaH*. All strains were grown in minimal medium containing 1 g/L 4HB as the sole source of carbon and energy.

## 4. Conclusions

We report here the first example of *E. coli* growing with PCA using the ortho-degradation pathway from *P. putida* KT2440. Chromosomal integration of the pathway allowed sustained growth in the absence of plasmids or antibiotics. Evolutionary optimization, resequencing, and reconstruction allowed the identification of pathway limitations. Increasing expression of this limiting enzyme increased the growth rate with PCA by roughly 2.5-fold, and allowed the further extension of the catabolic pathway to 4-HB. The optimized strain will be a useful platform for the characterization of aromatic catabolic pathways and for engineering conversion of aromatic compounds into value-added chemicals.

## 5. Acknowledgements

Genome resequencing and analysis was performed by Christa Pennacchio, Natasha Brown, Anna Lipzen, and Wendy Schackwitz at the Joint Genome Institute. The work conducted by the U.S. Department of Energy Joint Genome Institute, a DOE Office of Science User Facility, is supported by the Office of Science of the U.S. Department of Energy under Contract No. DE-AC02-05CH11231. Oak Ridge National Laboratory is managed by UT-Battelle, LLC, for the DOE under Contract No. DE-AC05-00OR22725.

## Author Contributions

SMC, DMK, JGE, and AMG designed, constructed and tested strains AG977 and AG978. JKM designed and performed experiments for evolutionary optimization, characterization, and pathway extension. SMC, AMG, and JKM wrote the paper.

## References

Blount, Z.D., Barrick, J.E., Davidson, C.J., Lenski, R.E., 2012. Genomic analysis of a key innovation in an experimental *Escherichia coli* population. Nature 489, 513–518. doi:10.1038/nature11514

Bugg, T.D.H., Ahmad, M., Hardiman, E.M., Rahmanpour, R., 2011. Pathways for degradation of lignin in bacteria and fungi. Nat. Prod. Rep. 28, 1883. doi:10.1039/c1np00042j

Chen, Y.-J., Liu, P., Nielsen, A.A.K., Brophy, J.A.N., Clancy, K., Peterson, T., Voigt, C.A., 2013. Characterization of 582 natural and synthetic terminators and quantification of their design constraints. Nat. Methods 10, 659–664. doi:10.1038/nmeth.2515

Chou, H.-H., Chiu, H.-C., Delaney, N.F., Segrè, D., Marx, C.J., 2011. Diminishing returns epistasis among beneficial mutations decelerates adaptation. Science 332, 1190–2. doi:10.1126/science.1203799

Clark, I.C., Melnyk, R.A., Youngblut, M.D., Carlson, H.K., Iavarone, A.T., Coates, J.D., 2015. Synthetic and Evolutionary Construction of a Chlorate-Reducing *Shewanella oneidensis* MR-1. MBio 6, e00282–15. doi:10.1128/mBio.00282-15

Datsenko, K.A., Wanner, B.L., 2000. One-step inactivation of chromosomal genes in *Escherichia coli* K-12 using PCR products. Proc. Natl. Acad. Sci. U. S. A. 97, 6640–5. doi:10.1073/pnas.120163297

Delaney, N.F., Kaczmarek, M.E., Ward, L.M., Swanson, P.K., Lee, M.-C., Marx, C.J., 2013. Development of an Optimized Medium, Strain and High-Throughput Culturing Methods for *Methylobacterium extorquens*. PLoS One 8, e62957. doi:10.1371/journal.pone.0062957

Department of Energy, U.S., 2016. 2016 Billion-Ton Report: Advancing Domestic Resources for a Thriving Bioeconomy, Volume 1: Economic Availability of Feedstocks. Oak Ridge, TN.

Diderichsen, B., 1980. *flu*, a metastable gene controlling surface properties of *Escherichia coli*. J. Bacteriol. 141, 858–67.

Doten, R.C., Ngai, K.L., Mitchell, D.J., Ornston, L.N., 1987. Cloning and genetic organization of the pca gene cluster from *Acinetobacter calcoaceticus*. J. Bacteriol. 169, 3168–74.

Espah Borujeni, A., Channarasappa, A.S., Salis, H.M., 2014. Translation rate is controlled by coupled trade-offs between site accessibility, selective RNA unfolding and sliding at upstream standby sites. Nucleic Acids Res. 42, 2646–59. doi:10.1093/nar/gkt1139

Harden, M.M., He, A., Creamer, K., Clark, M.W., Hamdallah, I., Martinez, K.A., Kresslein, R.L., Bush, S.P., Slonczewski, J.L., 2015. Acid-Adapted Strains of *Escherichia coli* K-12 Obtained by Experimental Evolution. Appl. Environ. Microbiol. 81, 1932–1941. doi:10.1128/AEM.03494-14

Harwood, C.S., Parales, R.E., 1996. The β-ketoadipate pathway and the biology of self-identity. Annu. Rev. Microbiol. 50, 553–590. doi:10.1146/annurev.micro.50.1.553

Herring, C.D., Raghunathan, A., Honisch, C., Patel, T., Applebee, M.K., Joyce, A.R., Albert, T.J., Blattner, F.R., van den Boom, D., Cantor, C.R., Palsson, B.Ø., 2006. Comparative genome sequencing of *Escherichia coli* allows observation of bacterial evolution on a laboratory timescale. Nat. Genet. 38, 1406–1412. doi:10.1038/ng1906

Hong, K.-K., Vongsangnak, W., Vemuri, G.N., Nielsen, J., 2011. Unravelling evolutionary strategies of yeast for improving galactose utilization through integrated systems level analysis. Proc. Natl. Acad. Sci. 108, 12179–12184. doi:10.1073/pnas.1103219108

Jiang, Y., Chen, B., Duan, C., Sun, B., Yang, J., Yang, S., 2015. Multigene Editing in the *Escherichia coli* Genome via the CRISPR-Cas9 System. Appl. Environ. Microbiol. 81, 2506–2514. doi:10.1128/AEM.04023-14

Jimenez, J.I., Minambres, B., Garcia, J.L., Diaz, E., 2002. Genomic analysis of the aromatic catabolic pathways from Pseudomonas putida KT2440. Environ. Microbiol. 4, 824–841. doi:10.1046/j.1462-2920.2002.00370.x

Kasai, D., Fujinami, T., Abe, T., Mase, K., Katayama, Y., Fukuda, M., Masai, E., 2009. Uncovering the Protocatechuate 2,3-Cleavage Pathway Genes. J. Bacteriol. 191, 6758–6768. doi: 10.1128/JB.00840-09

Kershner, J.P., McLoughlin, S.Y., Kim, J., Morgenthaler, A., Ebmeier, C.C., Old, W.M., Copley, S.D., 2016. A Synonymous Mutation Upstream of the Gene Encoding a Weak-Link Enzyme Causes an Ultrasensitive Response in Growth Rate. J. Bacteriol. 198, 2853–2863. doi: 10.1128/JB.00262-16

Kosuri, S., Goodman, D.B., Cambray, G., Mutalik, V.K., Gao, Y., Arkin, A.P., Endy, D., Church, G.M., 2013. Composability of regulatory sequences controlling transcription and translation in *Escherichia coli*. Proc. Natl. Acad. Sci. U. S. A. 110, 14024–9. doi:10.1073/pnas.1301301110

Linger, J.G., Vardon, D.R., Guarnieri, M.T., Karp, E.M., Hunsinger, G.B., Franden, M.A., Johnson, C.W., Chupka, G., Strathmann, T.J., Pienkos, P.T., Beckham, G.T., 2014. Lignin valorization through integrated biological funneling and chemical catalysis. Proc. Natl. Acad. Sci. U. S. A. 111, 12013–8. doi:10.1073/pnas.1410657111

Maruyama, K., Shibayama, T., Ichikawa, A., Sakou, Y., Yamada, S., Sugisaki, H., 2004. Cloning and Characterization of the Genes Encoding Enzymes for the Protocatechuate Meta-degradation Pathway of *Pseudomonas ochraceae NGJ1*. Biosci. Biotechnol. Biochem. 68, 1434–1441. doi: 10.1271/bbb.68.1434

Masai, E., Katayama, Y., Fukuda, M., 2007. Genetic and Biochemical Investigations on Bacterial Catabolic Pathways for Lignin-Derived Aromatic Compounds. Biosci. Biotechnol. Biochem. 71, 1–15. doi: 10.1271/bbb.60437

Michener, J.K., Camargo Neves, A.A., Vuilleumier, S., Bringel, F., Marx, C.J., 2014. Effective use of a horizontally-transferred pathway for dichloromethane catabolism requires posttransfer refinement. Elife 3. doi:10.7554/eLife.04279

Nichols, N.N., Harwood, C.S., 1997. PcaK, a High-Affinity Permease for the Aromatic Compounds 4-Hydroxybenzoate and Protocatechuate from *Pseudomonas putida*. J. Bacteriol. 179, 5056–5061.

Ornston, L.N., Stanier, R.Y., 1964. Mechanism of beta-ketoadipate formation by bacteria. Nature 204, 1279–83.

Ragauskas, A.J., Beckham, G.T., Biddy, M.J., Chandra, R., Chen, F., Davis, M.F., Davison, B.H., Dixon, R.A., Gilna, P., Keller, M., Langan, P., Naskar, A.K., Saddler, J.N., Tschaplinski, T.J., Tuskan, G.A., Wyman, C.E., 2014. Lignin valorization: improving lignin processing in the biorefinery. Science 344, 1246843. doi:10.1126/science.1246843

Salis, H.M., Mirsky, E.A., Voigt, C.A., 2009. Automated design of synthetic ribosome binding sites to control protein expression. Nat. Biotechnol. 27, 946–950. doi:10.1038/nbt.1568

Salvachúa, D., Karp, E.M., Nimlos, C.T., Vardon, D.R., Beckham, G.T., 2015. Towards lignin consolidated bioprocessing: simultaneous lignin depolymerization and product generation by bacteria. Green Chem. 17, 4951–4967. doi:10.1039/C5GC01165E

Schnetz, K., Rak, B., 1992. IS5: a mobile enhancer of transcription in *Escherichia coli*. Proc. Natl. Acad. Sci. U. S. A. 89, 1244–8.

Yona, A.H., Manor, Y.S., Herbst, R.H., Romano, G.H., Mitchell, A., Kupiec, M., Pilpel, Y., Dahan, O., 2012. Chromosomal duplication is a transient evolutionary solution to stress. Proc. Natl. Acad. Sci. 109, 21010–21015. doi:10.1073/pnas.1211150109

